# Time or distance: predictive coding of Hippocampal cells

**DOI:** 10.1101/2022.10.23.513401

**Authors:** Shai Abramson, Benjamin J. Kraus, John A. White, Michael E. Hasselmo, Genela Morris, Dori Derdikman

**Author notes:** Corresponding authors equally-contributing authors.

## Abstract

The discovery of place cells within the hippocampus has pointed to the importance of the hippocampus for navigation. The more recent discovery of hippocampal time cells has broadened the perspective of encoding in the hippocampus. An alternative hypothesis to the existence of time cells is based on the notion that hippocampal cells deduce location by integrating travelled distance (“path integration”). According to this alternate hypothesis, time cells, which fire at particular times when animals are running on a treadmill without changing location, actually encode accumulated distance on the treadmill. To examine this hypothesis, Kraus et al.^1^ performed treadmill experiments in which animals either ran for a fixed time or a fixed distance with varying velocities. Two distinct coding modes of hippocampal principal cells were found. Some cells encoded travelled distance and others elapsed time, thus refuting the notion that all hippocampal cells were performing path integration. Using the data from these experiments, we asked whether the two populations depended on the type of task the rats were engaged in. We show that the type of experiment determined the cells’ encoding, such that in fixed-distance experiments distance-encoding cells dominated, while on fixed-time experiments time-encoding cells dominated. These results suggest that the cells’ encoding contains a predictive element, dependent on the important variables of the experiment.

## Introduction

The hippocampus plays an important role in spatial processing and episodic memory ^2 3^. Spatial processing and navigation are supported by spatially tuned cells throughout the hippocampal formation, such as place cells within the hippocampus, which sparsely encode location within an environment ^4 5^. The subsequent discovery of time cells in the hippocampus ^1 6 7 8 9^, which encode time within an episode, suggests that these may contribute to the building blocks of episodic memory formation. The similarity of properties of time cells and place cells has led to a unifying concept of the hippocampus, as encoding dimensions required in order to organize relevant information. We thus asked whether the encoding of hippocampal neurons changes according to behavioral context and task demand. We used previously published data by Kraus et al.^1^, from an experiment which sought to resolve an inherent ambiguity in the interpretation of time cells. Time cells were initially reported in animals running on a running wheel without control of velocity^6^ (although time cells were also reported for stationary rats ^7^). This led to a potential ambiguity between encoding of time and of distance, due to the fact that, in fixed velocities, distance may be encoded by integration of time. Kraus et al.^1^ varied the velocity of rats running in place on a treadmill, and found subpopulations of hippocampal cells that encoded time, other cells that encoded path-integrated distance and additional cells that encoded both time and distance. These experiments were composed of two types of recording sessions. In one type of session, in all the trials the running duration remained constant at different velocities, whereas in the second type, the treadmill runs accumulated up to a constant distance, at different velocities. We hypothesized that in this experiment, the task demand (i.e. constant time vs. constant distance) determined the type of activity exhibited in the corresponding session. We re-analyzed the data according to the type of behavioral session and found a direct relation between the class of most active cells and the type of session in which they were recorded. In sessions in which the rats ran for a fixed time, the cells’ population was dominated by time encoding cells, while in sessions where they ran for a fixed distance, the population was dominated by distance encoding cells.

## Results

To examine the dependence of hippocampal coding on task demand, we analyzed data based on experiments by Kraus et al.^1^, which aimed to differentiate between cells encoding time and cells encoding distance in the hippocampus. In these experiments, six rats were trained to run on a treadmill in the central stem of a figure-8 maze (Figure 1a), with their noses at a water port, in order to “clamp” their behavior and location. In each session, consisting of 31-57 runs, the treadmill was operated either for a fixed time or for a fixed distance, where on each run the velocity was set to a speed randomly chosen in the range of 35-49 cm/sec (Figure 1b,c). The rats were forced to alternate their post-treadmill turns between right and left. Three of the six rats were trained and recorded exclusively in fixed-distance or fixed-time sessions, while the other three rats were trained and recorded in sessions of both types.

**Figure 1.**
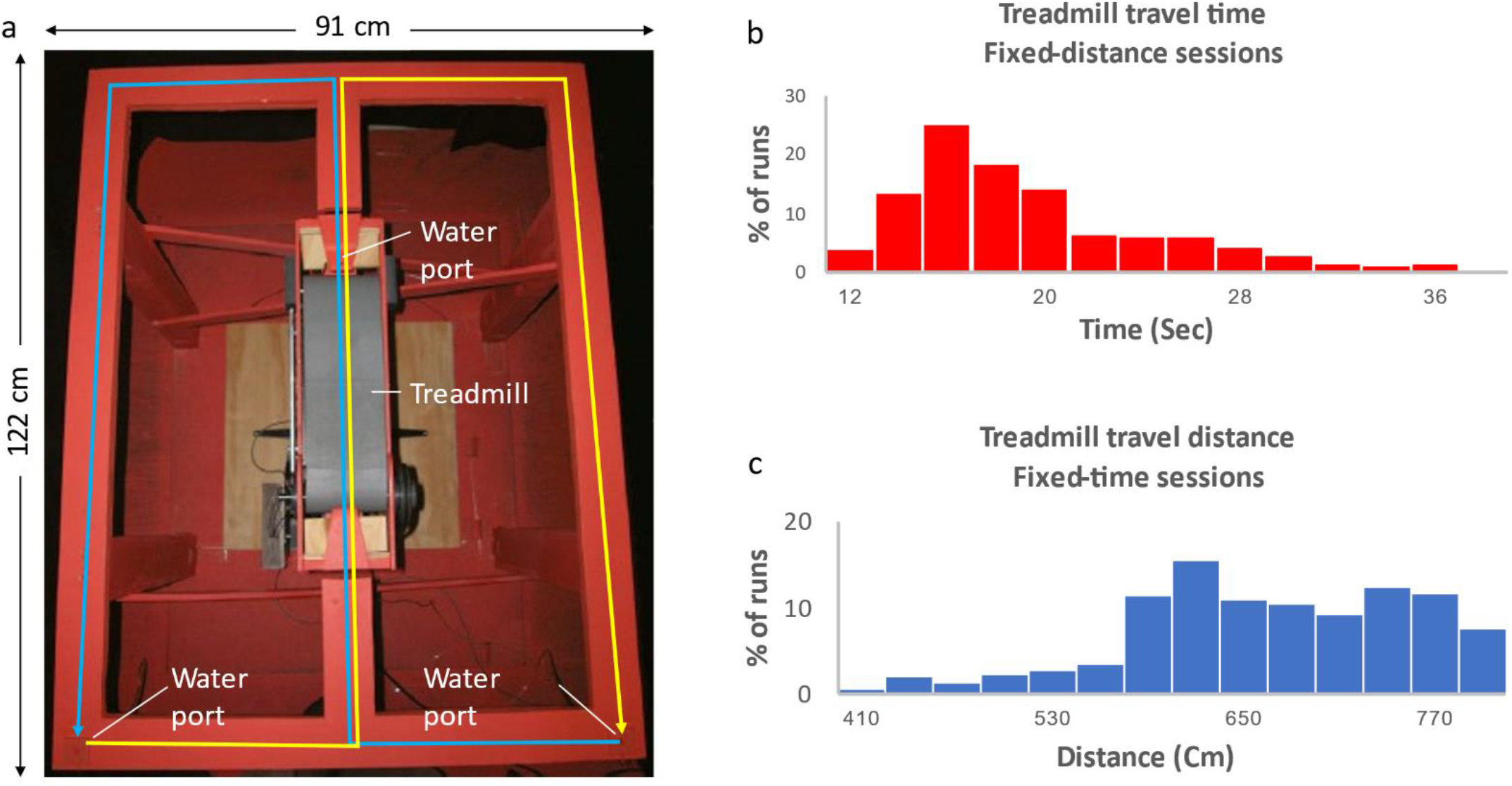
Experimental setup. **a** Picture of the figure-eight maze with treadmill (gray belt) located in the central stem. Water ports are located near the treadmill and at the two lower corners of the maze. Blue line indicates right-to-left alternation; yellow line indicates left-to- right alternation. Image taken from Kraus et al.^1^ **b,c** Distribution of the fixed-time sessions treadmill travel times (b) and the fixed-distance sessions treadmill travel distances (c).

Kraus et al. reported that some cells preferentially encoded the distance the rat had run on the treadmill while other cells preferentially encoded the time from the start of the treadmill movement. We hypothesized that the type of task employed in each session (i.e. fixed-time vs. fixed-distance) would determine the encoding of the neurons (i.e. time-based vs. distancebased). We therefore analyzed the cells on a run-by-run basis, as follows: For each neuron, we defined the onset of response in each run, to examine its relation to the treadmill’s velocity. We classified time-encoding cells as those, in which the onset time was almost independent of the treadmill velocity. We classified distance-encoding cells, on the other hand, as those, in which the onset time tended to be proportional to the treadmill velocity. To examine this classification, we determined the firing onset of each cell in each run and determined the cell’s properties according to three classifiers (see Methods section). Briefly, we determined the response onset time as the first bin with activity prior to the peak of firing. We defined a CellType index, based on the distance and time variances (see Analysis Methods), such that for a perfect time cell CellType=1 and for a perfect distance cell CellType=-1.

Of 930 cells recorded we analyzed 738 cells with at least 10 runs showing firing peaks greater than 0.5 Hz. Only cells with peak firing rates occurring during the treadmill run were included in the analysis.

As previously reported in Kraus et al., we observed both distance cells, demonstrating a firing peak at constant distances the animal traveled on the treadmill, and time cells, showing a firing peak at constant times from the treadmill start (Figure 2). In line with our hypothesis, there was a clear relation between the types of experiment and the distribution of time coding and distance coding neurons. In fixed-distance sessions, the neurons exhibited a significant majority (69%) of distance cells. By Contrast, in fixed-time sessions time encoding cells dominated (68%). Of the 465 neurons recorded in fixed-distance sessions, the CellType index classified 322 cells as distance cells and 143 as time cells (Figure 3a). Conversely, in the fixed-time sessions 87 of 273 neurons were classified by this index as distance cells and 186 of 273 were classified as time cells. The relation between the cell type and the experiment type is significant (*χ*^2^(1)=95.77, P<<0.001 for the total cells population). These proportions were maintained when classifying by other metrics (see supplementary Methods). In 5 out of 6 animals, there was a dependence between the cell’s encoding and the session type, with high significance (Figure 3-figure supplement 2). (*χ*^2^(1)>12.2, P<<0.001), except in one animal (*χ*^2^(1)=2.06, P=0.15)

**Figure 2.**
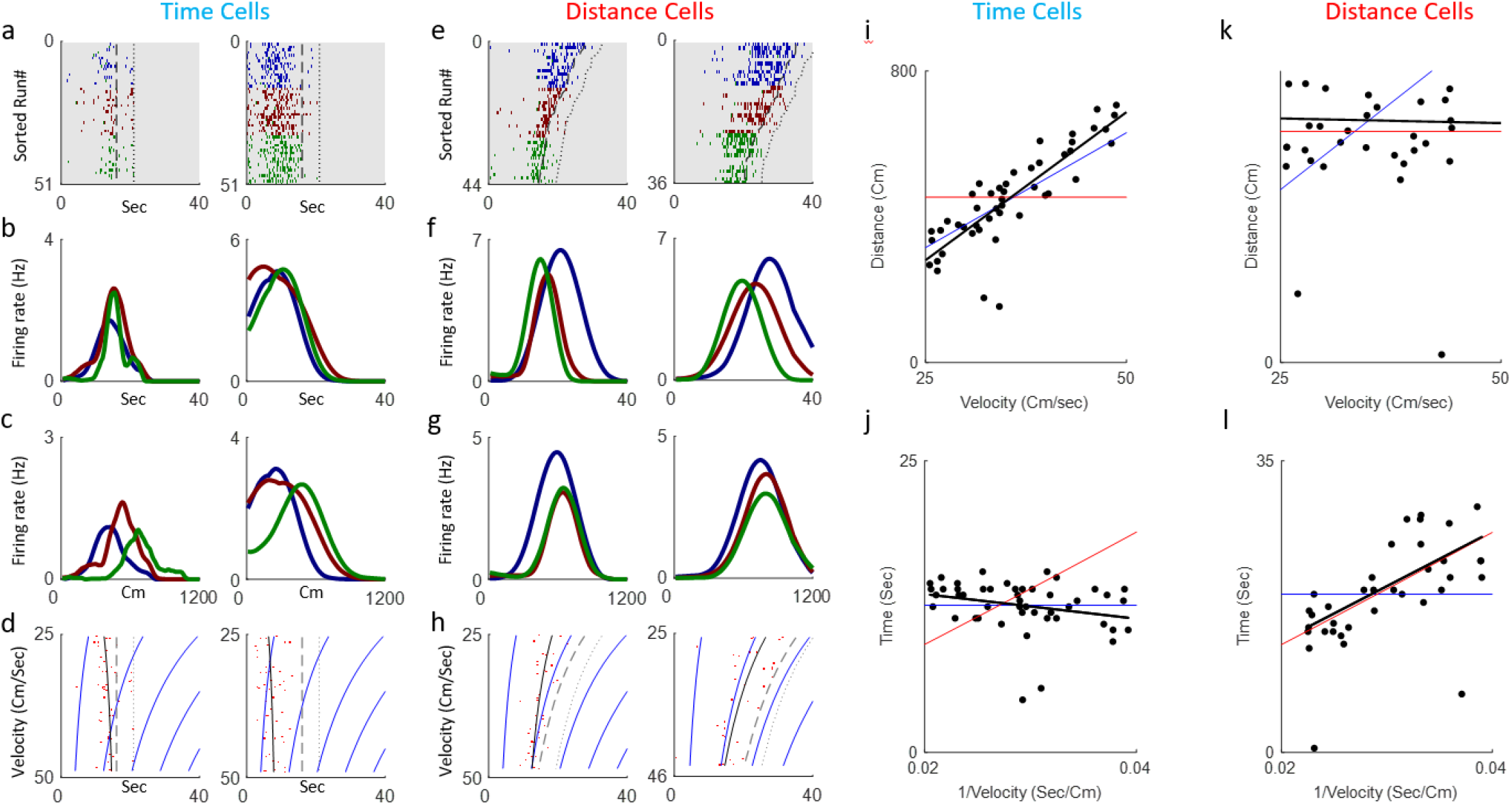
Distance and Time Cells coding. **a-h** Examples of two time-coding cells (columns 1-2) and two distance-coding cells (columns 3-4). Row a depicts neural firing as a function of the distance the animal traveled, sorted by the run’s velocities. The colors represent three velocity groups for which the tuning curves, by distance or time, are presented in rows b and c, respectively. Row d shows the onsets of each run (red dots) and their linear fit (black curve) to the relation 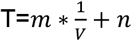 for the time cells and 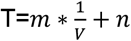 for the distance cells. The dashed curve represents the end-of-run time, while the dotted curve represents the end of the period analyzed (treadmill stop time, plus 5 seconds). The blue curves are equi-distance points in time. A black curve (the linear fit) which is parallel to the equidistance curves demonstrates a cell with strong distance coding. **i-l** Examples of the analysis of time (I,j) and time (k,l) encoding cells. Top row graphs depict distance vs. velocity and bottom row graphs depict Onset vs. 1/velocity. Red line represents a perfect Distance Cell, based on the average distance traveled until the onset time. Blue line represents a perfect Time Cell, based on the average time of the onset and the black line is the linear fit. The closer the black line slope is to the red line slope relative to the blue line slope, the more the cell is distance-encoding, while if the slope is closer to the blue line slope the cell is more time-encoding.

**Figure 3.**
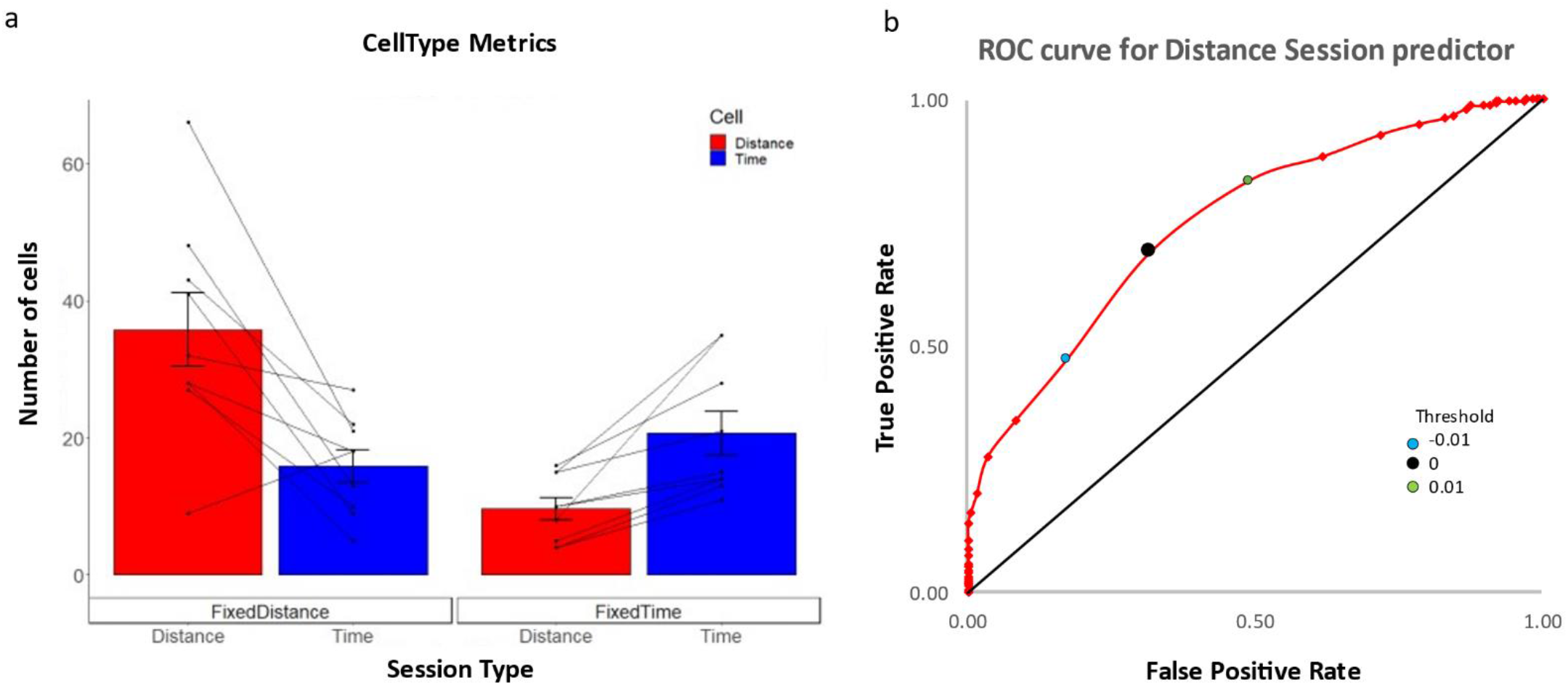
Distance and Time Cells classification. **a** Cells type classified by the CellType classifier, averaged over all animals and trials, for fixed-distance and fixed-time experiments. The whiskers show SEM, diagonal lines represent individual animals. **b** ROC curve (red) showing that the chosen discriminating threshold of 0 (black point) is optimal. The True Positive Rate (TPR) is the percentage of cells classified as distance cells on the fixed-distance session, while the False Positive Rate (FPR) is the percentage of cells classified as distance cells on the fixed-time sessions.

These results indicate that the dimension the cells encode (Time vs. Distance) is related to the session type (fixed time vs. fixed distance).

## Discussion

Classifying neuronal activity according to either time or distance dimensions revealed that the hippocampal population encoding strongly registers with the experiment type. In experiments where the treadmill running-time was fixed, the majority of cells encoded a given time from treadmill onset. In contrast, in experiments where the treadmill running-distance was fixed, the majority of cells encoded a specific accumulated distance from treadmill onset. It is worth noting that accumulated time in fixed-time experiments and accumulated distance in fixed-distance experiments may be used as predictors for the progress of the rat towards anticipated reward, which is given at the end of the treadmill run ^10 11^. As noted previously in Kraus et al.^1^ the same cells, which showed distance-encoding and time-encoding properties in the treadmill, were many times selective to places outside of the treadmill as well. To summarize, CA1 pyramidal cells can encode location, distance, or time, depending on the conditions of the experiment or task demand.

Consistency with task demands has been repeatedly demonstrated in hippocampal recording for diverse parameter spaces, such as auditory linear frequency ^12^, social mapping ^13 14^ or more abstract spaces ^15 16^. How task-relevant encoding is achieved? The activity of place cells and grid cells is commonly modeled using continuous attractor networks ^17 18 19 20 21 22 23 24 25^. Such networks may serve as a natural substrate for amplification of encoding of certain task dimensions, at the expense of others.

One possible mechanism for acquiring representations that are consistent with task structure involves an associative learning process. Such learning would modify all connections that were active in a particular trial but would only consistently modify those in which activity was invariant throughout the experiment, while other connections associated with inconsistent representations will average out. Thus, in time-fixed experiments, those connections that are consistent with time would be strengthened, while in distance-fixed experiments those that are consistent with distance would gain strength. Reward or task-completion could generate a prolonged signal, which, upon coinciding with cells that receive time and distance signals, will strengthen the synaptic connections of the contributing signal. Consequently, those cells will gradually encode either distance or time, depending on the type of experiment (Figure 4).

**Figure 4.**
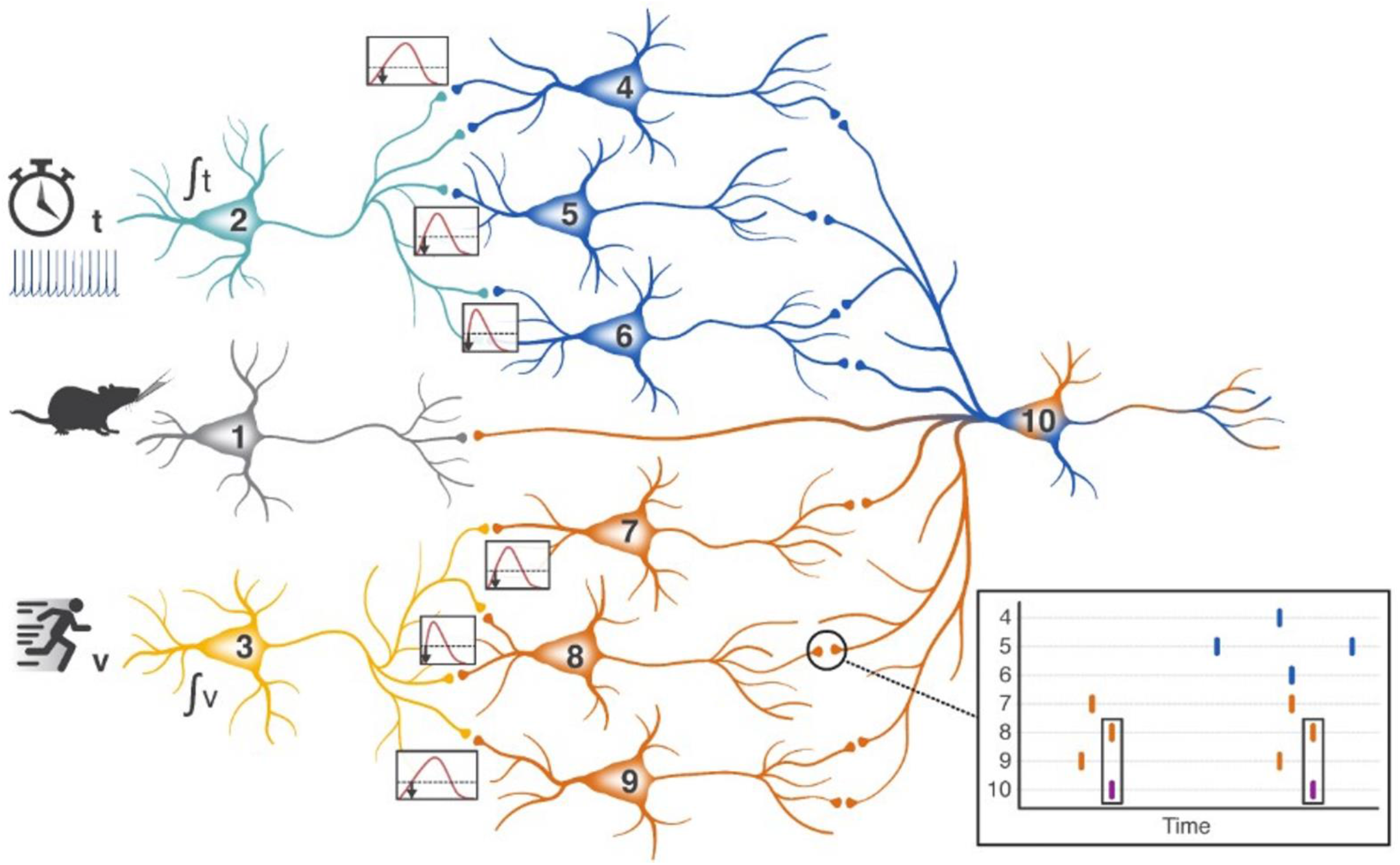
Predictive Coding model illustration. Cell 2 (possibly from an upstream brain region) performs a temporal integration function from treadmill’s start time, and connects to cells 4 to 6, which fire at different times per their synaptic potential, consequently encoding diverse periods of time. Cell 3 receives the animal’s speed (possibly from the MEC), integrating it and connecting to cells 7 to 9, which fire at different thresholds and accordingly encode diverse distances. Cell 10 receives the outputs of cells 4 to 9, as well as a reward or treadmill stop prediction signal from cell 1. Initially, the reward input dominates, causing cell 10 to fire at the reward prediction time. When the reward prediction coincides with the firing of any cell from 4 to 9 (cell 8 in the inset example), the connection from the coinciding cell is strengthened through Hebbian learning. Accordingly, after several trials within a given experimental day, cells best encoding the prediction will dominate and generate an output signal at cell 10, regardless of the reward.

Irrespective of the exact mechanism explaining the results of this study, the hippocampus is adaptive in its cells’ encoding and seems to be capable to tune them to the parameters best describing the task.

## Analysis Methods

We used the data provided by Kraus et al ^1^, containing the neurons firing times, the treadmill movement times and the treadmill velocity. The data was analyzed using custom Matlab scripts.

We divided the treadmill moving times into 200 ms time bins (other bin resolutions between 100 msec and 500 msec were tested and provided similar results). For each neuron, in each run, the response onset was defined as the first bin with activity prior to the peak in firing. For each neuron, the onsets of all runs within the same session were analyzed. For a perfect time encoding cell, we expect the onset times *T_i_* to be independent of the treadmill velocity *V_i_*, whereas for a perfect distance cell, we expect the onsets to be linearly dependent on the reciprocal velocity (equation 1). Therefore, for a time encoding cell, the product of the fit coefficient *k* and velocity would be small compared to the offset coefficient *q,* while for a distance encoding cell the fit coefficient would approximate the estimated encoded distance.

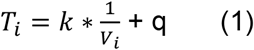

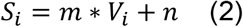

Similarly, we analyzed the distance the animal traveled on the treadmill *S_i_* until the onset versus the treadmill velocity *V_i_*. Following the relation in equation 2, a perfect distance encoding cell will have a product of the linear coefficient m and the velocity small relative to the offset coefficient n, while time encoding cell will have an estimated encoded time of *m.*

The CellType classifier was based on the variances of the onsets and the distances related to those onsets:

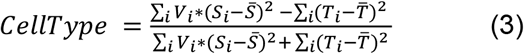

CellType is in the range of −1 to 1. For a perfect time encoding cell, the onset variance 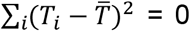 and hence CellType=1. For a perfect distance encoding cell, the distance variance (multiplied by the respective velocity in order to match units) 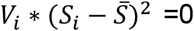 and hence CellType=−1.

To classify the cell as either a time or a distance encoding cell, we have chosen a discriminating threshold using the Receiver Operating Characteristic graph (Figure 2d), which plots the True Positive Rate defined as the percentage of cell classified as distance cells in the fixed-distance session, against the False Positive Rate defined as the percentage of cells classified as distance cells in the fixed-timed sessions.

We found that the optimal threshold was 0. This threshold classifies 69% of the cells as Distance in the Distance sessions while only 32% on the Time sessions. Accordingly, we classified a cell as a time-cell if CellType>0 and as a distance-cell if CellType<0.

To ensure the activity peak is not missed, we have extended the analysis to 5 seconds past the treadmill stop time. Otherwise, if a cell activity is concentrated towards the treadmill stop, the calculated onset may be influenced by the truncated activity time and show a false relation of the cell type activity vs the time or distance. Moreover, since the truncated data time relates to the experiment type, whether time-fixed or distance-fixed, this could create a false bias of such a relation.

The relation between the type of cell classified in the above metrics and the session type was then tested through Pearson’s chi-squared test using two categories (DOF=1). The expected distribution of the cells was calculated based on the total number of cells, of each type, out of total cells number, in all sessions. The null hypothesis was defined as no dependency of the cells type distribution on the session type (either fixed-time or fixed-distance). On the per-animal analysis, for animals that were recorded only at a single type of session, we used the distribution of the cell types according to their distribution in all animals’ cells population.

## Supplementary Methods

Additional metrics defined and used for classifying the cells encoding:

The “FIT” classifier is defined as follows:

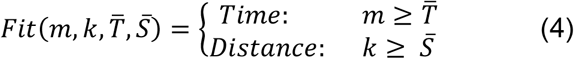

Where *m* and *k* are the linear fit slope coefficients (from equations 1 & 2), 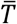 is the average firing onset time and 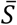 is the average distance the animal traveled until the onset.

A stricter classifier based on this measure, tested the statistical significance of the linearity in equations 1 and 2, through F-statistics. We classified a cell as distance encoding if the null hypothesis that there is no linear relation between the distance and velocity was rejected with p<0.05. We classified a cell as time encoding if the null hypothesis that there is no linear relation between the onset and the reciprocal velocity was rejected with p<0.05.

Results using these classifiers are shown in Figure 3-figure supplement 1

**Figure 2-figure supplement 1.**
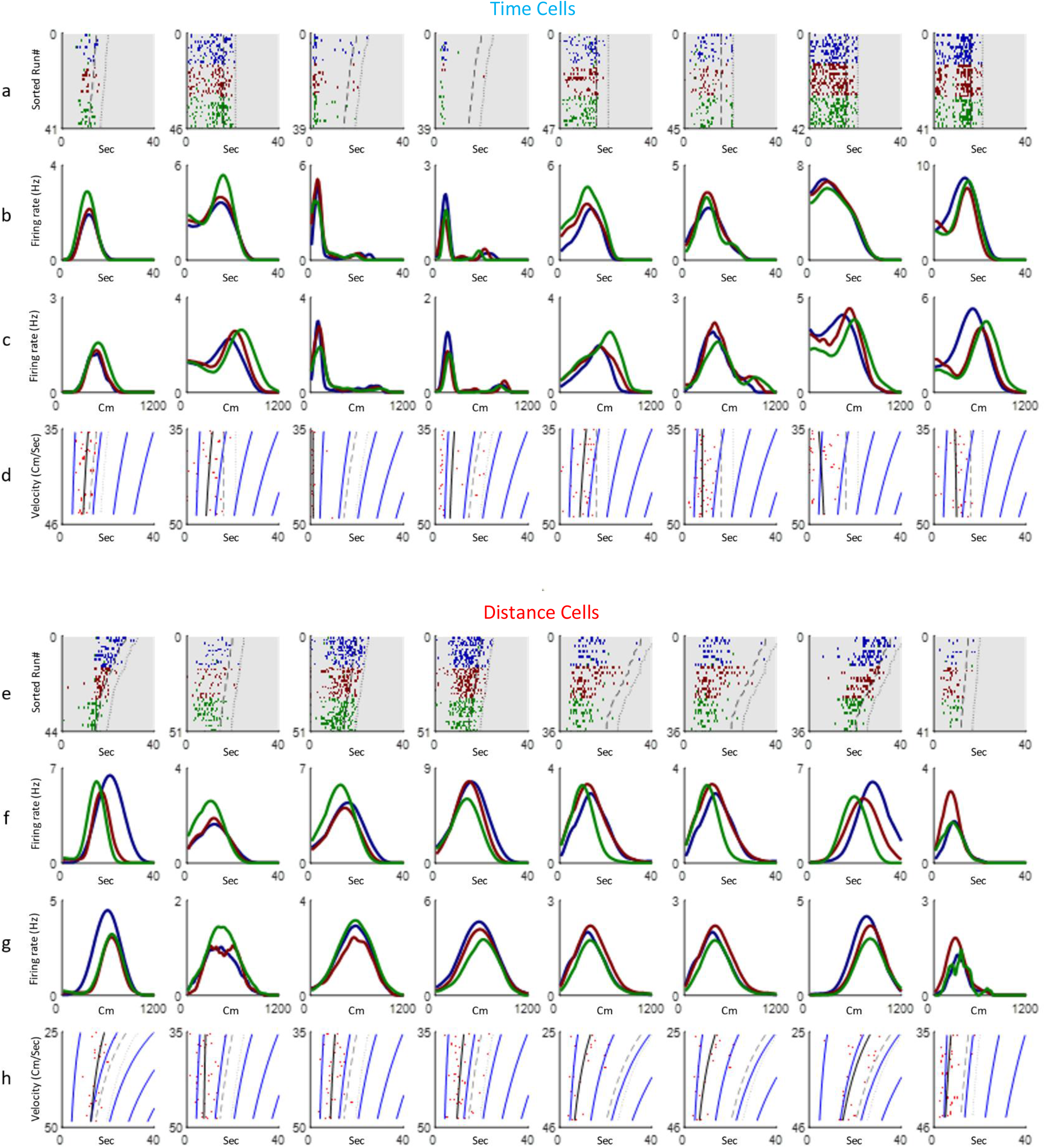
Additional examples of cells showing time coding (a-d) and distance encoding (e-h). Rows a,e depict neural firing as a function of time, sorted by the run’s velocities. The colors represent three velocity groups for which the tuning curves, either by time or distance, are presented in rows b,f and c,g respectively. Rows d,h show the onsets of each run (red dots) and their linear fit (black curve) based on the relation 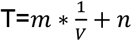 for the time cells and 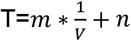 for the distance cells. Dashed curves represent treadmill stop time and dotted curves represent end of period analyzed (treadmill stop time plus 5 seconds). Blue curves are equi-distance points in time while dashed curve represents end of run time. The more a linear fit curve is vertical the more significant is the time coding of that cell.

**Figure 3-figure supplement 1.**
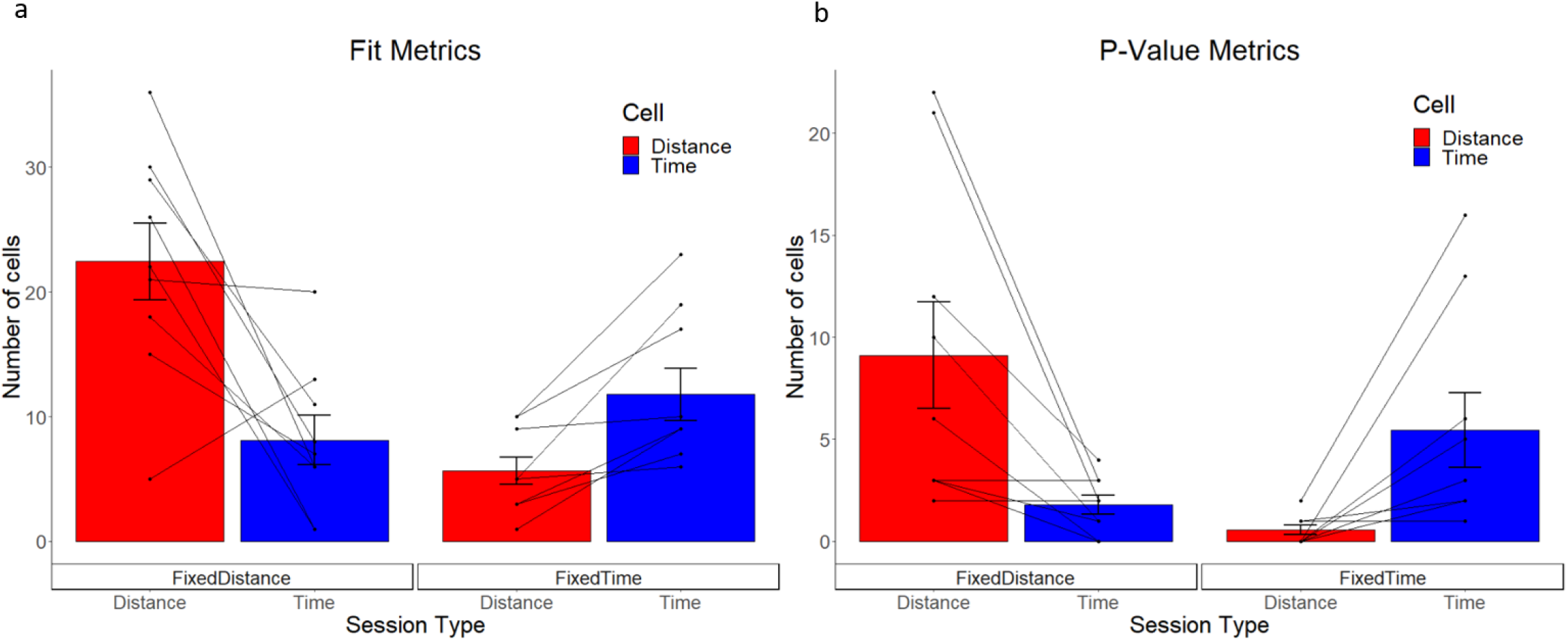
Cells type classified by the FIT metrics **(a)** and P-Value metrics **(b)**, averaged over all animals and trials, for fixed distance and fixed time experiments. The whiskers show the SEM and the points and the lines connecting the points represent individual animal results.

**Figure 3-figure supplement 2.**
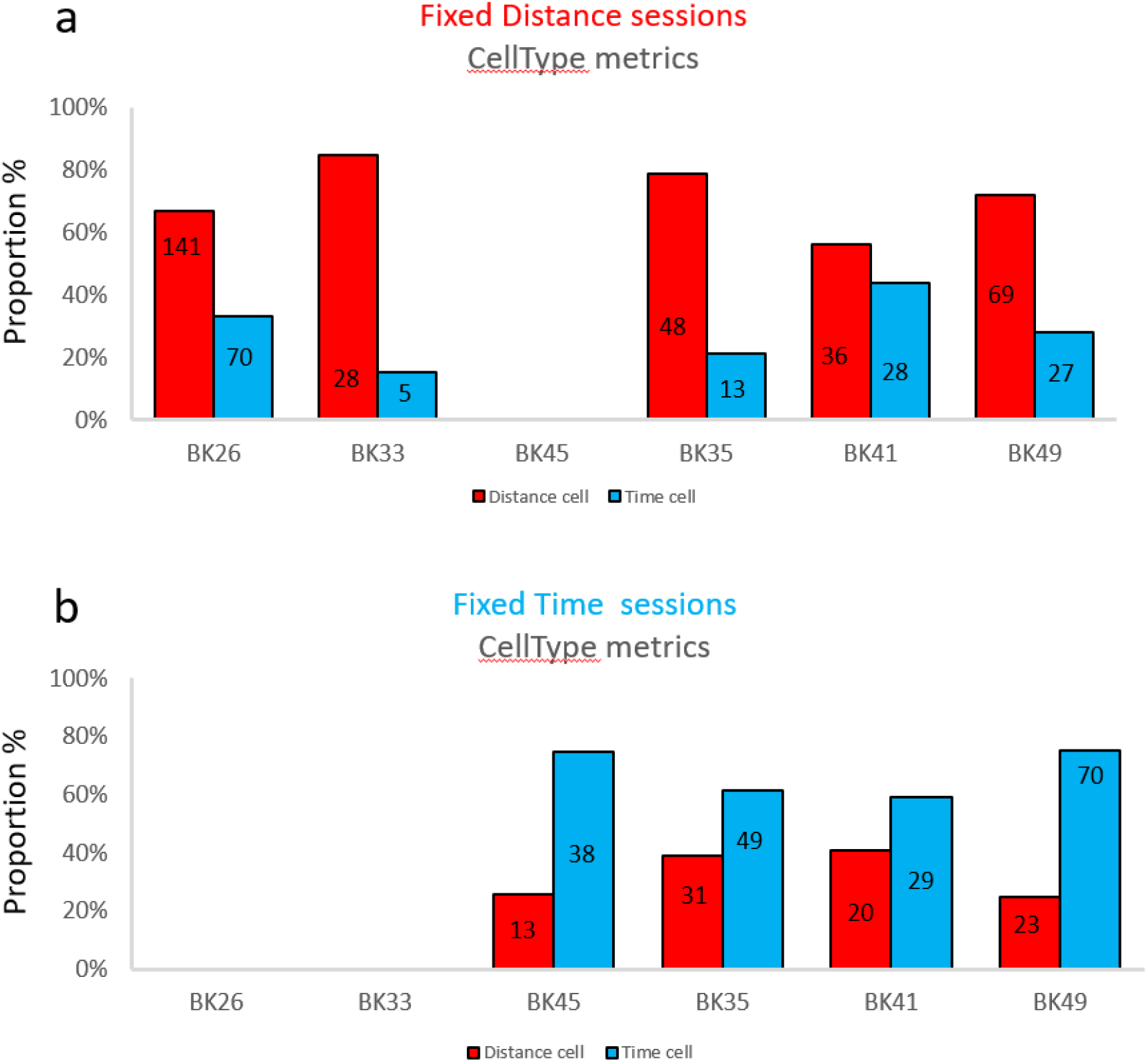
The distribution of Time and Place cells, per animal, at various classifier metrics, for Time and Distance sessions. Animals BK26, BK33 participated only in distance sessions, while BK45 participated only in a time session, and BK35, BK41 and BK49 participated in both types of sessions. All distributions show a significant dependency on the session type (χ^2^(1)>12.2, P<<0.001), except for animal BK41 (χ^2^(1)=2.06, p=0.15). The bars represent the percentage of the respective cell type from the entire population of cells in the respective session. The numbers inside the bars represent the number of cells. For the single session animals, the reference distribution used for the χ^2^ test was taken from the entire population distribution (46% Distance Cells and 54% Time Cells).

## Data availability

Matlab and R scripts are available on https://github.com/derdikman/Abramson_code. Data, originally curated in ref. 1 and used in the current paper, is available on Dryad: https://doi.org/10.5061/dryad.ngf1vhhxp

## Acknowledgements

We thank Dafna Antes for help in preparation of Figure 4. This research was supported by the Israel Science Foundation (grants Nos. 2655/18 and 2183/21), by the German-Israeli Foundation (GIF I-1477- 421.13/2018), by a grant from the US-Israel Binational Science Foundation (NIMH-BSF CRCNS BSF:2019807, NIMH:R01 MH125544-01 to DD) by A Rappaport Institute Collaborative research grant and by the Prince Center for the Aging Brain.

